# Single-cell transcriptome analysis of Physcomitrella leaf cells during reprogramming using microcapillary manipulation

**DOI:** 10.1101/463448

**Authors:** Minoru Kubo, Tomoaki Nishiyama, Yosuke Tamada, Ryosuke Sano, Masaki Ishikawa, Takashi Murata, Akihiro Imai, Daniel Lang, Taku Demura, Ralf Reski, Mitsuyasu Hasebe

## Abstract

**Background:** Next-generation sequencing technologies have made it possible to carry out transcriptome analysis at the single-cell level. Single-cell RNA-sequencing (scRNA-seq) data provide insights into cellular dynamics, including intercellular heterogeneity as well as inter- and intra-cellular fluctuations in gene expression that cannot be studied using populations of cells. The utilization of scRNA-seq is, however, restricted to specific types of cells that can be isolated from their original tissues, and it can be difficult to obtain precise positional information for these cells *in situ*.

**Results:** Here, we established single cell-digital gene expression (1cell-DGE), a method of scRNA-seq that uses micromanipulation to extract the contents of individual living cells in intact tissue while recording their positional information. Furthermore, we employed a unique molecular identifier to reduce amplification bias in the cDNA libraries. With 1cell-DGE, we could detect differentially expressed genes (DEGs) during the reprogramming of leaf cells into stem cells in excised tissues of the moss *Physcomitrella patens*, identifying 6,382 DEGs between cells at 0 h and 24 h after excision. We found substantial variations in both the transcript levels of previously reported reprogramming factors and the overall expression profiles between cells, which appeared to be related to their different reprogramming abilities or the estimated states of the cells according to the pseudotime based on the transcript profiles.

**Conclusions:** We developed 1cell-DGE with microcapillary manipulation, a technique that can be used to analyze the gene expression of individual cells without detaching them from their tightly associated tissues, enabling us to retain positional information and investigate cell–cell interactions.

## Background

Whole plants can be regenerated from tissue samples such as branch cuttings or detached leaves via a callus [1], a mass of undifferentiated cells able to initiate shoot- and root-stem cells in the presence of the appropriate phytohormones [2]. Several genes have been shown to function in regenerating cells [3, 4]; however, the elucidation of the transcriptome profiles involved in the regeneration process of each cell is a major challenge as it is not currently possible to separate and identify the limited numbers of stem cells that randomly emerge in a callus during regeneration.

A number of single-cell RNA-sequencing (scRNA-seq) methods utilizing next generation sequencing (NGS) have been developed to prepare cDNA libraries from isolated single cells containing trace amounts of RNA [5, 6]. Hundreds to thousands of single isolated cells derived from the human fetal cortex or mouse retinas have been simultaneously prepared into sequencing libraries using automated single-cell preparation systems such as Fluidigm C1 [7] and inDrop [8, 9]. By assessing the heterogeneity of expression profiles between individual cells in a population, including rare cell types, biological events such as different cell cycle stages and transcription bursts have been identified, revealing the trajectories of developmental cell states that were not previously detectable in transcriptome analyses of samples containing multiple cells [10, 11]. In plants, single-cell transcriptome analyses of root cells in the flowering plant *Arabidopsis thaliana* (Arabidopsis) revealed a transition of cell identity during root regeneration [12-14]. scRNA-seq has great potential for providing new biological insights into regeneration; however, using the methods described above, the positional information of the cells within their tissue is lost during the isolation process. Furthermore, it can be difficult to detach single cells from the tissues and organs of many plant species because their cell walls consisting of carbohydrate and proteoglycan polymers strongly adhere to each other.

The moss *Physcomitrella patens* (Physcomitrella) is a basal land plant with a simple body plan, including leaves formed of a single cell layer [15], which facilitates its observation and manipulation at the cellular level [16, 17]. When a Physcomitrella leaf is cut, some of the cells facing the cut change into chloronema apical stem cells without the addition of exogenous plant hormones, enabling the entire moss body to be regenerated [18]. Several genes involved in this reprogramming have been characterized. Cyclin-dependent kinase A (PpCDKA) and cyclin D (PpCYCD;1) regulate the reentry into the cell cycle [18]. The *WUSCHEL-related homeobox 13* (*PpWOX13*) genes are upregulated during reprogramming and required for the tip growth characteristic of the chloronema apical stem cells [19]. The Cold-Shock Domain Protein 1 (PpCSP1) and PpCSP2, orthologous to the mammalian reprogramming factor Lin28A, also positively regulate reprogramming in Physcomitrella [20]. Furthermore, a transcriptome analysis of whole excised leaves during reprogramming revealed that the expression levels of more than 3,900 genes were altered within 24 hours after excision [21].

When Physcomitrella leaves are excised, only some of the leaf cells facing the cut are reprogrammed, while other cells neighboring the cut, as well as the intact cells that do not face the cut, are not reprogrammed [18]. It is therefore difficult to distinguish between genes specifically expressed in the reprogramming cells and those expressed in non-reprogramming cells. Understanding the *in situ* regulation of reprogramming in an excised leaf is a challenge; when two neighboring leaf cells are isolated together, only one is reprogrammed, even though almost all cells isolated on their own can autonomously reprogram into protonema apical cells [22]. This suggests the presence of cell–cell interactions between neighboring cells during reprogramming; however, the molecules and genes responsible for this mechanism have not been identified, partially because of the difficulty in isolating a single cell to investigate its transcriptome during the reprogramming process. When a pair of adjacent cells are isolated, both show features of the early phases of reprogramming, such as nuclear expansion and the expression of cell cycle-related genes; however, these become diminished in the non-reprogrammed cell [22]. This suggests that the reprogrammed cells not only inhibit reprogramming in their neighbors, but that they actively revert their neighboring cells back to a leaf cell state. Although this is a good model for studying cell–cell interactions during reprogramming, it has meant that the mechanisms by which stem cells are determined and the factors involved in the inhibitory effect of the reprogrammed cells on their neighbors are poorly understood.

To explore the genes involved in cell–cell interactions of reprogramming in Physcomitrella leaves, we established a single cell transcriptome analysis method using microcapillary manipulation to physically extract the contents of individual living cells within a tissue and prepare a cDNA library of their trace amounts of RNA. We also introduced a unique molecular identifier (UMI) [23] to the cDNAs to reduce the amplification bias when using PCR.

## Results

### Extraction of the contents of single cells in excised leaves

We employed microcapillary manipulation to isolate the contents of individual leaf cells in Physcomitrella while recording their positional information. Our initial attempts to generate cDNA from the extracted contents of entire cells were rarely successful, presumably because the central vacuole occupies ~ 90% of the plant cell volume [24] and accumulates RNases that degrade RNA molecules [25]. Since the transcriptomes of the isolated nuclei are reported to be similar to those of the whole cells [26, 27], we extracted cell contents including nuclei labelled with a fusion protein (NGG) composed of a nuclear-localizing signal [28], sGFP (synthetic green fluorescent protein) [29], and GUS (β-glucuronidase) [30] under an estrogen-inducible system [18, 31].

We excised the distal half of Physcomitrella leaves and, after 24 hours, sucked the nucleus and surrounding cytoplasm from individual leaf cells facing the cut (Figure 1, Additional file 1: Supplementary Movie S1). We synthesized cDNA from the RNA in the cellular contents without any purification, and amplification using PCR, a quantitative PCR (qPCR) was used to determine the transcript levels of four genes: *NGG*, *CYCLIN D;1* (*PpCYCD;1*), *ELONGATION FACTOR 1α* (*PpEF1α*), and *TUBULIN α1* (*PpTUA1*) (Additional file 2: Supplementary Figure S1). The transcript levels of *PpCYCD;1* (sample standard deviation: s = 7.9423) and *PpTUA1* (s = 7.9431), which are known to be upregulated during reprogramming [12, 32], varied among the contents of the different cells, while those of the positive controls *NGG* (s = 2.8616) and *PpEF1α* (s = 1.9492) were mostly stable in all contents, as expected. The variation in the *PpCYCD;1* and *PpTUA1* transcripts suggests that the isolated single cells include those in different stages of reprogramming as well as those not undergoing reprogramming, as previously observed [18]. These results indicate that cDNAs derived from single-cell contents extracted using a microcapillary can be used to detect gene expression in single leaf cells of Physcomitrella, as was previously shown for isolated cell contents of individual cells of tobacco, potato, and cucumber [32].

**Figure 1.**
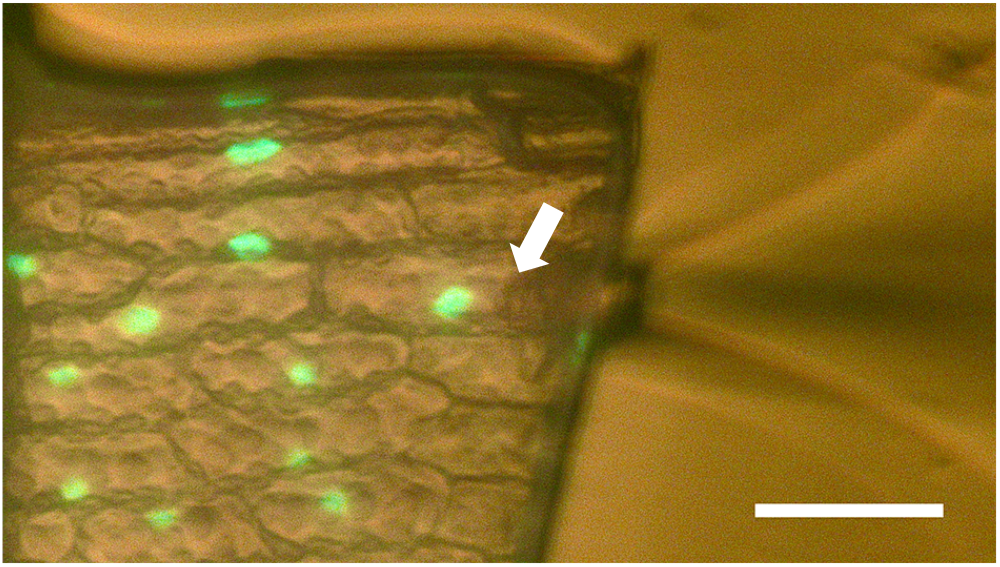
Extraction of the contents of an individual cell using microcapillary manipulation. An arrow indicates the tip of a microcapillary extracting the contents of a cell facing the cut edge of an excised Physcomitrella leaf expressing GFP in its nuclei. Imaged using GFP fluorescence under dim bright-field illumination. Scale bar: 50 μm.

### Preparation of cDNA libraries from traces of total RNA

To make a cDNA library of at least 40 femtomoles (2.4×10^10^ molecules) for sequencing on Illumina next-generation sequencers, we amplified the single-cell cDNAs using PCR. Two methods of attaching an adaptor to the 3′ ends of the first-strand cDNAs were evaluated: template switching [33] and polydeoxyadenines (dA) tailing [34-36]. Template switching introduces several non-templated deoxycytosine residues to the 3′ ends of the first-strand cDNA before synthesizing the complementary strands. Poly(dA) tailing extends the poly(dA) region at the 3′ ends of the first-strand cDNAs using a terminal deoxynucleotidyl transferase, followed by the annealing of a specific primer with 20-nt oligo (dT) sequences to the poly(dA) tail. After second-strand synthesis using both types of cDNAs, we performed qPCR to detect cDNA quality and quantity using various RNA amounts for the libraries. Using the poly(dA) tailing libraries, the decreases in the crossing point PCR cycle (Cp) values of *PpTUA1* cDNA were proportional to increases in the total RNA amounts from 10 pg to 10 ng (Additional file 2: Supplementary Figure S2). By contrast, the changes in the Cp values when using the template switching libraries did not demonstrate this relationship. We therefore adopted poly(dA) tailing for our single-cell transcriptome analysis.

Another problem is the amplification bias in cDNAs when using PCR. These biases are caused by differences in amplification efficiencies, which depend on the length, nucleotide contents, and sequences of the DNA fragments, as well as stochastic fluctuations [37]. To reduce the template-dependent biases, we adopted the unique molecular identifier (UMI) method, in which random barcode sequences are introduced into the first-strand cDNA at the time of reverse transcription [23]. When the sequence reads are mapped to the reference genome [38, 39], reads with the same UMI are considered to have originated from the same cDNA. To test the UMI method, we sequenced the cDNA libraries derived from 5 μg and 20 pg of total RNA extracted from protonema cells from Physcomitrella (Additional file 2: Supplementary Figure S3, Additional file 3: Supplementary Table S1). In the two 5-μg library replicates, 780,202 and 972,450 reads were mapped onto the Physcomitrella v3.3 gene model, which were unified to 680,993 and 832,593 UMI counts, respectively. In the two 20-pg library replicates, 606,512 and 660,397 reads were mapped, and were unified to 82,229 and 101,533 UMI counts, respectively. We found that the read counts of each gene in the 5-μg libraries were strongly correlated between the duplicated samples (R^2^ = 0.9891), even if the UMIs were not unified (R^2^ = 0.9743). On the other hand, the read counts in the 20-pg libraries tended to vary if the UMIs were not unified (R^2^ = 0.8610); however, the UMI counts of each gene were strongly correlated between duplicated samples, to a similar level as those of the 5-μg libraries (R^2^ = 0.9677). We found similar tendencies for replicates of the External RNA Controls Consortium (ERCC) RNA spike-in mix, which were added to the total RNA samples as an external control (Additional file 2: Supplementary Figure S3c, d) [40]. Furthermore, the UMI counts reflected the nominal concentrations of the ERCC RNA spike-in mix, although the correlation was somewhat lower in the 20-pg libraries than the 5-μg libraries (Additional file 2: Supplementary Figure S3e–h).

### Preparation of cDNA libraries from the contents of single cells and NGS

To apply the methods described above to individual Physcomitrella leaf cells, we made cDNA libraries from single cells facing the cut of an excised leaf after 0 h and 24 h (Additional file 2: Supplementary Figure S4). The cell contents, including the nuclear region marked by the NGG, were extracted using microcapillary manipulation from 32 cells at 0 h and 34 cells at 24 h after the leaves were excised. The content of each cell was transferred to a PCR tube and cDNA was synthesized using reverse transcription with an exonuclease I treatment, poly(dA) tailing, second-strand synthesis, and cDNA amplification. Subsequently, the cDNA quality and quantity were measured, and the samples that showed a peak of cDNAs between 500 bp and 5,000 bp in length were purified by removing the byproducts detected between 100 bp and 400 bp. To prepare the Illumina sequencing libraries, four or five samples with different multiplex sequences were mixed equally in a single batch, and were subsequently subjected to fragmentation, end-repair, dA-tailing, adaptor-ligation, library enrichment, and library purification.

After quantification and qualification, these batches of NGS libraries were equally mixed and sequenced on an Illumina HiSeq sequencer set to generate 126-bp single-end reads and 18-bp index reads, including 8 bp of multiplex index and 10 bp of UMI. A total of 384,993,923 reads were obtained and cleaned by trimming and filtering to remove inaccurate sequences and ribosomal sequences, respectively. As a result, 98.1% (377,495,213) of the reads were sorted to their respective samples using the multiplex index sequences, generating 2.8 million to 8.5 million reads per sample for 66 samples (Additional file 2: Supplementary Figure S5, Additional file 3: Supplementary Table S2). Mapping these reads to Physcomitrella gene models, means of 5,566,262 and 4,966,189 mapped reads were obtained for the samples taken at 0 h and 24 h, respectively, which equated to mapping rates of 90.9% and 92.6%, respectively. These mapped reads were unified to remove duplicated reads located at the same gene locus and with the same UMI sequences, because the UMI sequences were introduced during the reverse transcription step and later amplified using PCR. Only sequences with different UMI sequences at the same gene locus were therefore considered as distinct cDNAs in the quantification of the original numbers of transcripts. For the cells sampled at 0 h and 24 h, the mean UMI counts were 102,145 and 91,851, respectively, and the UMI-unified rates, indicating the rate of duplicated reads (same locus with the same UMI), were 98.3% and 98.3%, respectively (Additional file 2: Supplementary Figure S6, Additional file 3: Supplementary Table S2). At 0 h and 24 h, the mean numbers of transcribed genes per sample were 5,277 and 7,297, respectively.

Generally, validating transcriptome data involves comparing the transcript levels of internal control genes with a similar expression level among all samples; however, it was difficult to choose an appropriate gene because the expression of many genes, including those which are generally accepted as housekeeping genes (e.g., *GLYCERALDEHYDE-3-PHOSPHATE DEHYDROGENASE* and *α-TUBULIN*), were found to fluctuate substantially at the single-cell level (Additional file 2: Supplementary Figure S1). We therefore checked the quality of the 1cell-DGE data using a statistically analyzed population of single-cell transcriptome data in the SinQC package [41], judging the outliers based on the statistics of mapping rates, the number of detected genes, and read complexity. All but one sample (31 cells at 0 h and 34 cells at 24 h after leaf excision) passed this evaluation with a max false positive rate (FPR) of 0.05 and using the following settings: TPM Cutoff: 1, Spearman’s test p-value: < 0.001, Pearson’s test p-value: < 0.001 (Additional file 3: Supplementary Table S2). These data were therefore used for further analyses.

To estimate how many reads are adequate for single-cell profiling using 1cell-DGE, we calculated the number of detected genes and UMI-unified rates within a limited number of reads randomly extracted from 1cell-DGE data at 0 h and 24 h (Additional file 2: Supplementary Figure S7). We did not detect any significant differences in the tendencies of these statistics between the 0-h and 24-h samples or among the selected index sequences. Although the numbers of detected genes increased as the number of sampled reads increased, the rate of change slowed as more sampled reads were considered. The UMI-unified rates also increased as the number of sampled reads increased, although they appeared to have close to an asymptotic relationship. At 2 million and 5 million reads, UMI-unified rates of 98.0% and 98.4%, respectively, were calculated for the samples taken at 0 h. For the samples taken after 24 h, UMI-unified rates of 97.1% and 98.0%, respectively, were calculated.

### Expression profiles of individual cells at 0 h and 24 h after leaf excision

To detect differentially expressed genes (DEGs) in the 1cell-DGE data taken at 0 h and 24 h after leaf excision, we carried out a statistical analysis after normalization using the iterative differentially expressed gene exclusion strategy (iDEGES) method [42]. A total of 6,382 genes were identified as DEGs, of which 2,382 and 4,000 genes were expressed at higher levels in the samples taken at 0 h (0 h-high) and 24 h (24 h-high), respectively, when calculated using the criterion of a false discovery rate (FDR) < 0.01. Using these gene expression profiles, we performed a hierarchal clustering and found that profiles for 0 h and 24 h were clearly categorized into separate populations, indicating characteristic transcript profiles (Figure 2).

**Figure 2.**
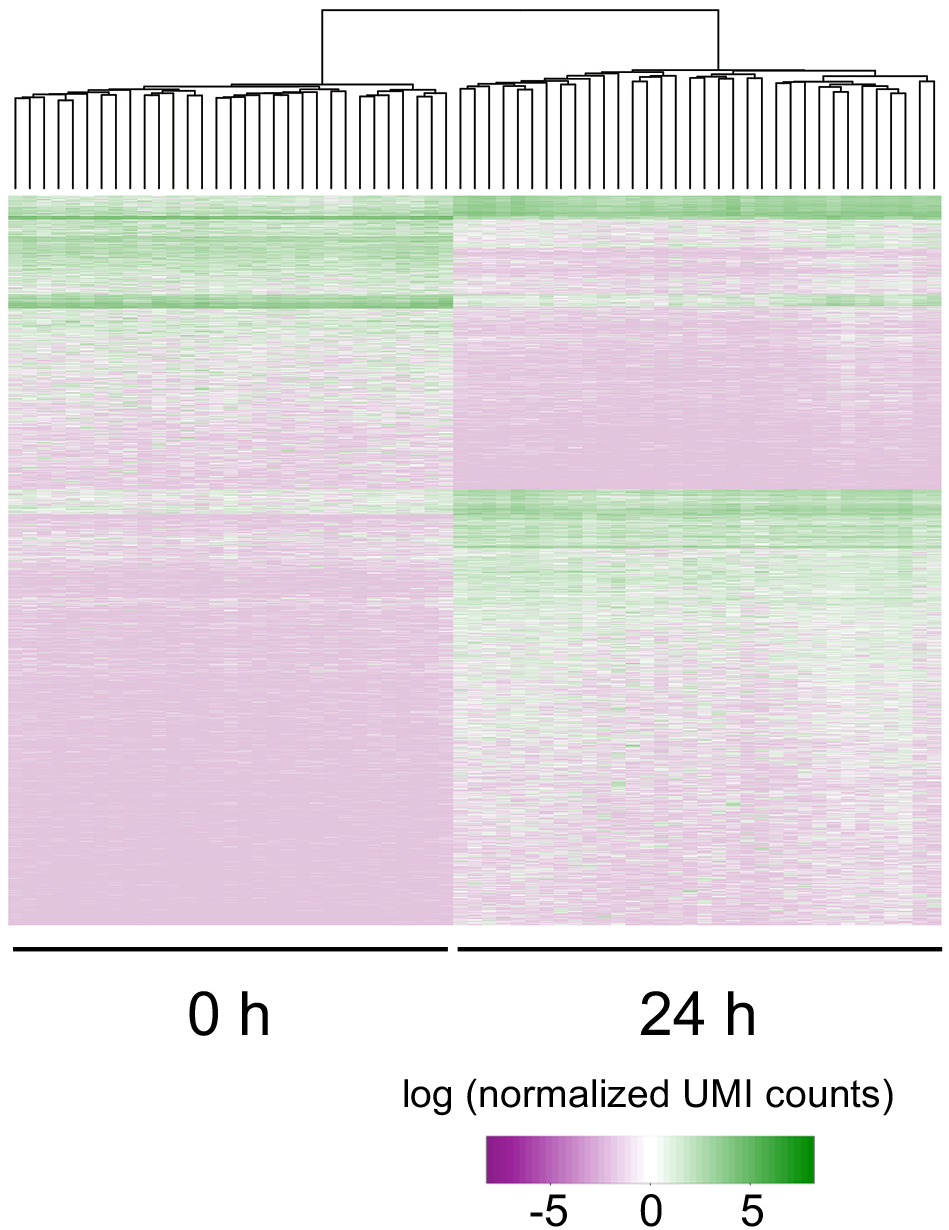
Hierarchal clustering of differentially expressed genes (DEGs) in individual cells at 0 h and 24 h after leaf excision. Using 1cell-DGE data taken from 31 and 34 cells at 0 h and 24 h after leaf excision, respectively, 6,382 genes were identified as DEGs (FDR < 0.01) after normalization with iDEGES using the TCC package [42]. Of these, 2,382 and 4,000 were more highly expressed in the cells at 0 h and 24 h, respectively. Hierarchal clustering was performed using hclust in the stats package. The colored bars indicate the normalized UMI counts as the expression levels of the DEGs on a log scale.

We performed a gene ontology (GO) term enrichment analysis for the 1,978 of the total 2,382 DEGs at 0 h and 3,648 of the total 4,000 DEGs at 24 h that were putatively homologous to annotated Arabidopsis genes (Figure 3). Using GOSlim_plants to categorize the genes, we revealed an enrichment of genes involved in the responses to stress and abiotic and biotic stimuli, the generation of precursor metabolites and energy, metabolic processes involving cellular amino acids and their derivatives, lipid metabolic processes, catabolic processes, post-embryonic development, reproduction, and cellular transport in both the 0 h-high and 24 h-high DEGs. In addition, the GO terms of photosynthesis, secondary metabolic process, and response to endogenous/external stimulus were enriched at 0 h, whereas those of cell growth, cell cycle, cell differentiation, embryonic development, DNA and protein metabolic process, biosynthetic process, translation, carbohydrate metabolic process, anatomical structure morphogenesis, and cellular component organization were enriched at 24 h.

**Figure 3.**
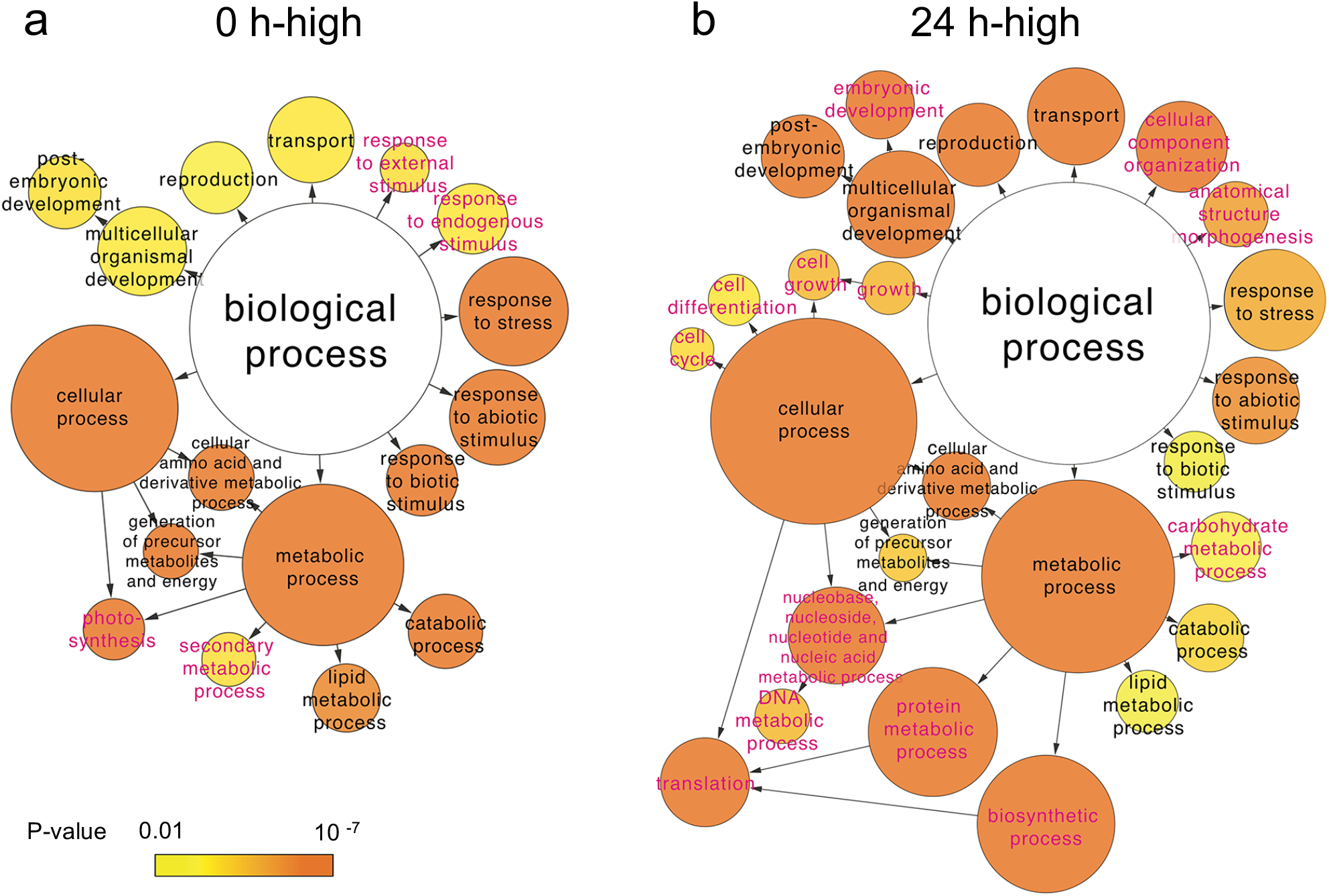
GO term enrichment analysis of differentially expressed genes (DEGs) in individual cells at 0 h and 24 h after leaf excision. A total of 1,978 of the 2,382 DEGs more highly expressed at 0 h and 3,648 of the 4,000 DEGs more highly expressed at 24 h after leaf excision could be annotated based on their homology to genes in *Arabidopsis thaliana*. Using the associated Arabidopsis gene identifiers, a GO term enrichment analysis was performed using cytoscape v.3.4.0 with the BinGO plug-in, and the ontology of biological processes was assessed using GOSlim_plants. The terms in magenta text indicate sub-categories are only represented in the DEGs more highly expressed at either 0 h or 24 h after leaf excision. The circles are colored based on the statistical significance of their enrichment.

In previous studies, transcriptome analyses of whole excised leaves during reprogramming were performed using 5′DGE [19, 21]. We therefore compared the DEGs identified using 1cell-DGE with those reported using the 5′DGE method for whole excised leaves (Figure 4). After remapping the 5′DGE data onto the Physcomitrella v3.3 gene models [39] and counting the read tags for each gene locus, 2,578 and 651 DEGs with a FDR < 0.01 were detected as 0 h-high and 24 h-high genes, respectively. A total of 751 of the 0 h-high DEGs were commonly identified in both the 1cell-DGE and 5′DGE analyses, while 395 of the 24 h-high DEGs were commonly identified between datasets.

**Figure 4.**
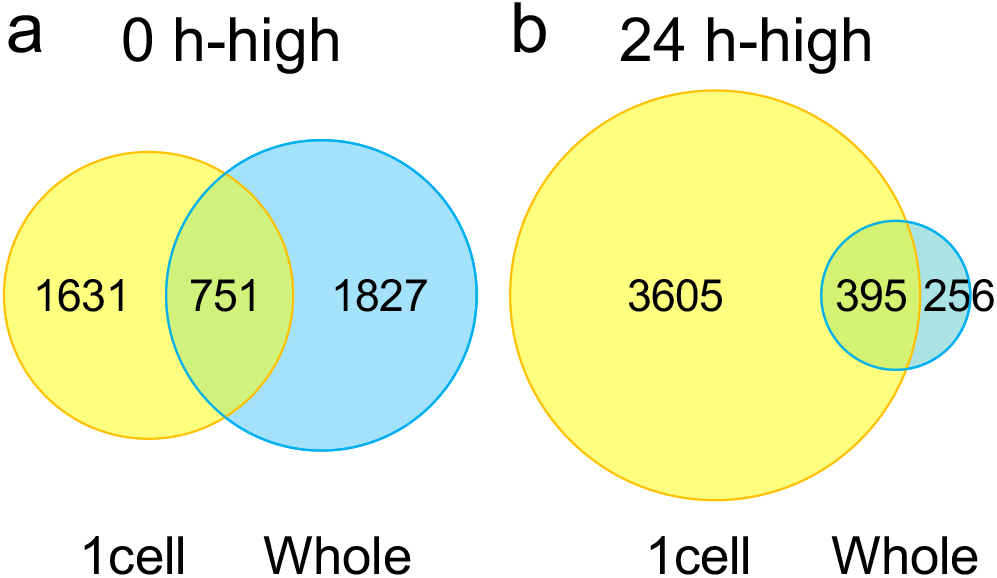
Comparison of differentially expressed genes (DEGs) between 1cell-DGE and whole leaf-5’DGE. Venn diagrams show the number of DEGs identified using 1cell-DGE and 5′DGE in single cells of excised leaves or the whole leaf, respectively, which were more highly expressed at 0 h or 24 h after leaf excision. The 5′DGE data from [19, 21] were remapped onto Physcomitrella v3.3 gene models [39] and normalized using the iDEGES method with the TCC package [42].

We then checked the expression levels of the top 10 DEGs detected by the statistical significance of their differences at 0 h and 24 h using a q-value (Figure 5). Pp3c23_13700 (unknown), Pp3c1_21540 (aluminium induced protein-like), Pp3c4_7680 (membrane protein, putative), Pp3c4_7130 (unknown), Pp3c13_7000 (glyoxal oxidase-related protein-like), Pp3c4_26000 (chaperone DnaJ-domain superfamily protein-like), Pp3c16_16490 (unknown), Pp3c5_25650 (unknown), Pp3c9_7780 (calcium-dependent lipid-binding family protein-like), and Pp3c4_30240 (TOXICOS EN LEVADURA 2-like) were selected as the top 10 DEGs more highly expressed at 0 h. Pp3c13_5750 (lactoylglutathione lyase / glyoxalase I family protein-like), Pp3c10_4900 (unknown), Pp3c4_29000 (unknown), Pp3c26_150 (unknown), Pp3c10_4280 (bHLH protein), Pp3c3_10800 (adenosine kinase 2-like), Pp3c15_7380 (dihydrodipicolinate reductase-like), Pp3c12_4560 (expansin A9-like), Pp3c14_8260 (succinyl-CoA ligase, alpha subunit-like), and Pp3c1_11820 (unknown) were selected as the top 10 DEGs more highly expressed at 24 h.

**Figure 5.**
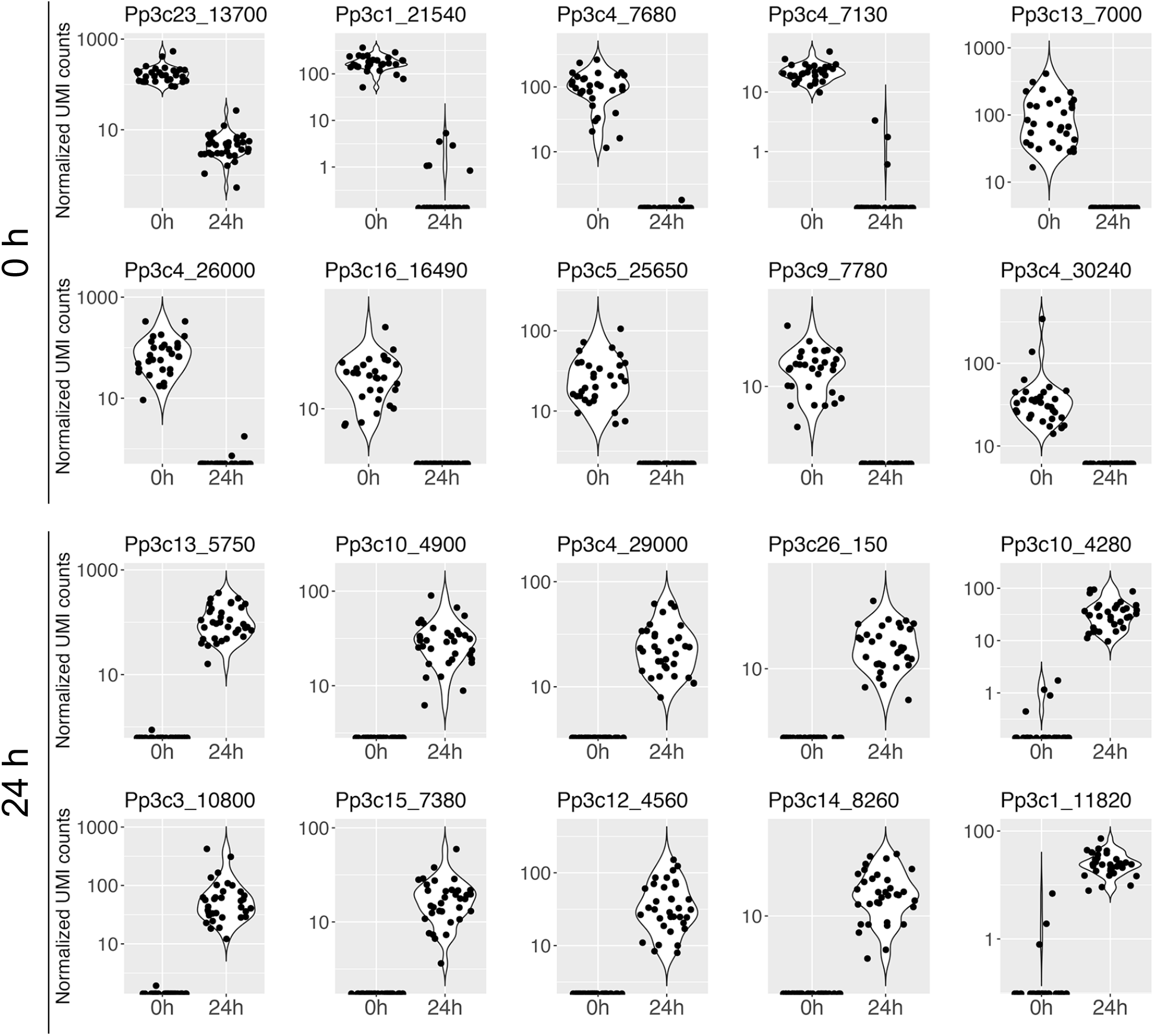
Expression profiles of 20 genes differentially expressed in single cells at 0 h and 24 h after leaf excision. The expression profiles of the genes which ranked in the top 10 DEGs at 0 h (upper two rows) and 24 h (bottom two rows) are shown as violin and jitter plots. The top 10 DEGs were selected by ranking the DEGs based on their m-values, p-values, and q-values calculated with the TCC package [42].

In order to take full advantage of single-cell transcriptome data, it is possible to calculate the pseudotime, a hypothetical time scale estimating the transition between cell states during development and differentiation based on similar gene expression profiles [11, 43]. First, an independent component analysis (ICA) was carried out to reduce the dimensions of the gene expression profiles (Figure 6). Like the hierarchal clustering, nodes indicating the individual samples in the ICA were clearly separated between cells sampled at 0 h or 24 h after excision (Figure 6a). Furthermore, we could not find any relationship between the cell profiles for the extracted nuclear condition, leaf excision date, cDNA amount, or byproduct contamination (Additional file 2: Supplementary Figure S8), further confirming the correlation between the ICA result and the other criteria. When each point in the ICA plot was colored according to its pseudotime, almost all points for both the 0-h and 24-h samples were found to be arranged in order of their pseudotime (Figure 6b).

**Figure 6.**
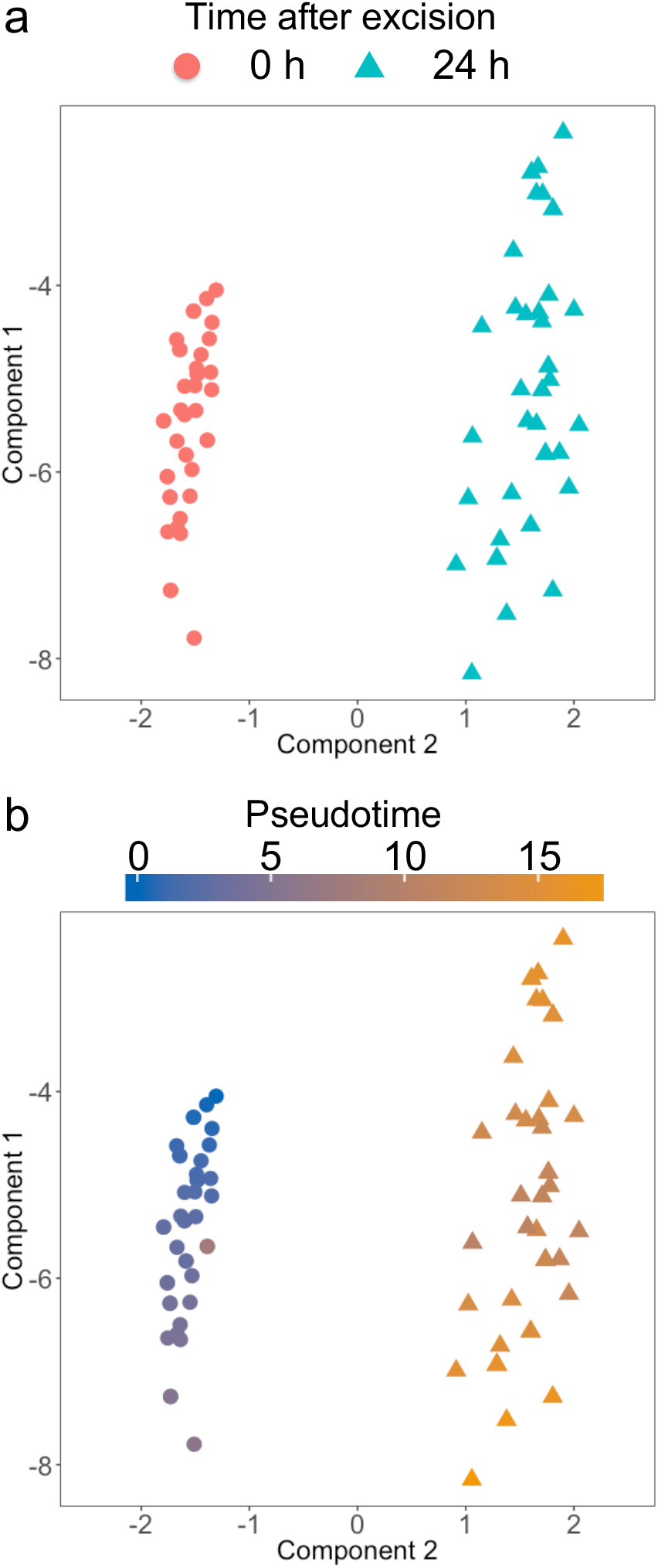
Independent component analyses of the 1cell-DGE data. Normalized 1cell-DGE data were calculated to reduce the dimensions of the expression profiles using an independent component analysis, and were plotted with the monocle package [43]. Each dot indicates an independent cell sample categorized by the time it was sampled after excision (a) or the pseudotime (b).

When the expression profiles of the Physcomitrella reprogramming-related genes *PpCSP1*, *PpCSP2*, and *PpCYCD;1* [18, 20] were plotted against pseudotime (Figure 7), they were generally found to be expressed at low levels in the early phase of pseudotime, with the exception of several cells with high *PpCSP1* expression. Further along the pseudotime scale, *PpCSP1* was the most highly expressed in cells at 24 h after the leaf excision. In contrast, *PpCYCD;1* expression varied substantially among cells at 24 h after the leaf excision, which is likely attributable to the heterogeneity in the reprogramming ability of the cells at the cut edge [22].

**Figure 7.**
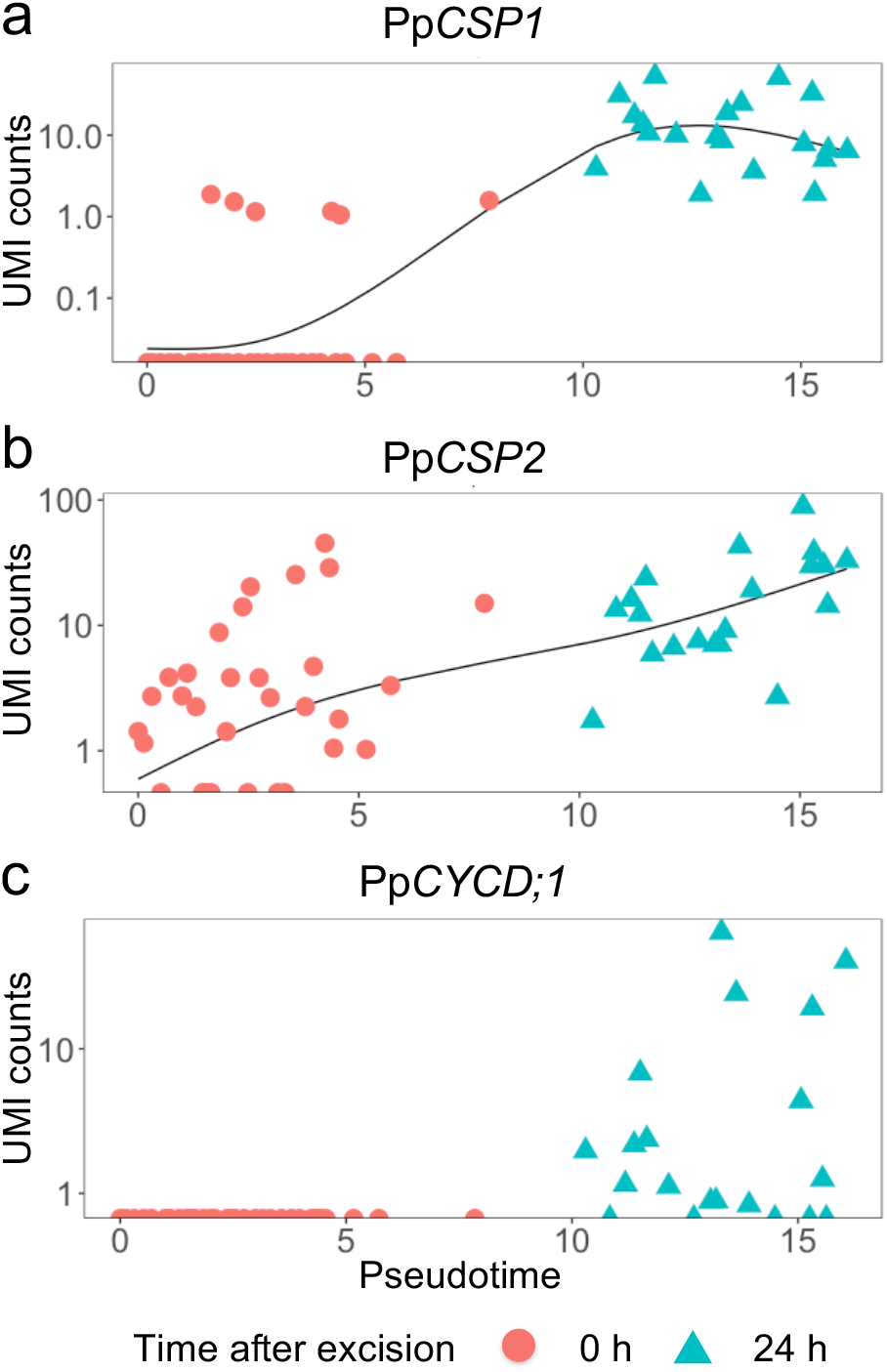
Expression profiles of reprogramming-related genes ordering by pseudotime. *PpCSP1* and *PpCSP2*, genes encoding cold-shock protein 1 and 2 [20]; *PpCYCD;1*: gene encoding cyclin D;1 [18]. These were calculated and plotted using the monocle package [43]. Each dot indicates an independent cell depicted as the time it was sampled after the excision.

We also compared the correlation measures between the pseudotime and NGS statistics. Using Hoeffding’s *D* test of independence for nonparametric and non-monotonic relationships [44], we identified a low correlation between the mapped read counts and the pseudotime (*D* = 0.014, p = 0.0493), but found a moderate correlation between the number of detected genes and the pseudotime (*D* = 0.285, p = 10^−8^) (Additional file 2: Supplementary Figure S9).

## Discussion

### Single cell transcriptome analysis using microcapillary manipulation

Since the first report of scRNA-seq [34], many methods of scRNA-seq have been developed and improved [5, 6, 45], but two major concerns have arisen: how can a single cell or its contents be isolated, and how can cDNA be efficiently and accurately prepared from trace amounts of RNA. The former challenge has been partly overcome using cell sorting with fluorescence-activated cell sorting (FACS) and microfluidics [7-9]; however, applying these systems require the separation of individual cells from their tissues or cultures. When the cells are separated using these techniques, the resulting samples no longer have accurate positional information. In addition, plant cells are often tightly attached to each other by their cell walls, making it difficult to mechanically or enzymatically detach them from each other while keeping their cellular contents intact. Microcapillary manipulation or laser microdissection can be used to extract the contents of single cells from tissues without detaching them, overcoming these challenges and enabling the transcriptomic analysis of individual cells while retaining positional information.

Reverse transcription polymerase chain reaction (RT-PCR) has previously been successfully performed with cDNA derived from the contents of single cells extracted from plant leaf cells using glass microcapillaries [32]; therefore, we attempted to prepare cDNA libraries from the contents of individual leaf cells from the moss Physcomitrella (Additional file 2: Supplementary Figure S4). Marking the nuclei with GFP (Figure 1, Additional file 1: Supplementary Movie S1) enabled us to reproducibly recover the cell contents including nuclear region using microcapillary manipulation and prepare cDNA that could be used for qPCR (Additional file 2: Supplementary Figure S1). Previous transcriptome analyses of isolated nuclei have demonstrated that they have similar expression profiles to those of whole cells [26, 27], indicating that this is an appropriate technique to use for the preparation of cDNA from individual cells.

Another issue was how to prepare cDNA from a small amount of RNA without excessive amplification bias which depends on the length, nucleotide contents, and sequences of the cDNAs [37]. Using conventional methods, NGS libraries are prepared for RNA-seq by purifying the mRNAs and fragmenting them before reverse transcription. In contrast, for scRNA-seq, the mRNAs are not purified, and are instead directly reverse-transcribed to cDNA from the crude cell contents. Generally, template switching or poly(dA) tailing is utilized to attach the adaptor oligonucleotides at the 3′ end of the cDNA after reverse transcription. In this study, we employed the latter technique for the 1cell-DGE based on our results (Additional file 2: Supplementary Figure S2). While we found template switching to be less effective than poly(dA) tailing, this could be improved by the use of a short template-switching oligo and locked nucleic acid (LNA)-linked nucleotides [46-48], and might be suitable for use with 1cell-DGE following such improvements.

The trace amount of first-strand cDNA generated from the RNA of single cells necessitates their amplification before they can be sequenced. To overcome an amplification bias, we introduced sequences of 6 or more random nucleotides, UMIs, to the cDNAs to enable their later discrimination (Additional file 2: Supplementary Figure S4) [23]. We designed 103 species of oligo (dT) nucleotides comprising 10 nt of UMI and 8 nt of multiplex index, which enabled us to identify the original index even if one substitution error occurred on the index sequence (Additional file 3: Supplementary Table S3). Using these RT oligos, we can mix samples with different multiplex indexes after the synthesis of the first-strand cDNA and subsequently prepare the NGS libraries as bulk samples. Moreover, as sequencing generated single-end reads of 50 bp to 126 bp with 18 bp of index reads, this approach is expected to reduce sequencing costs and more efficiently generate analyzable reads than conventional scRNA-seq with pair-end reads. After NGS, the original numbers of first-strand cDNAs can be estimated by unifying the reads derived from the same molecule, which are defined as the reads mapped to the same gene locus that possess the same UMI.

To test this, we performed pilot sequencing using total RNA purified from Physcomitrella protonema tissues. In 5-μg samples of cDNA, which had not been amplified using PCR, we found a similar determination coefficient (R^2^) between the read counts and the UMI counts (Additional file 2: Supplementary Figure S3a). By contrast, in the 20-pg samples, the R^2^ value of the UMI counts was higher than that of the read counts (Additional file 2: Supplementary Figure S3b). The quantification values of the ERCC RNA spike-in mix added to the pilot sequencing samples (Additional file 2: Supplementary Figure S3c, d) also showed high R^2^ values for both the read counts and the UMI counts between replicates. We confirmed that 1cell-DGE using UMIs enabled the highly reproducible quantification of cDNA from trace samples of RNA. In addition, we found a high correlation between the concentrations and UMI counts of the ERCC RNA spike-in mix in both the 5-μg and 20-pg samples (Additional file 2: Supplementary Figure S3e–h). The lower R^2^ values in the 20-pg samples may originate from the low coverage of the pilot sequencing (Additional file 3: Supplementary Table S1), and the drop-out reads known to be a feature of scRNA-seq [49, 50]. Our results therefore demonstrate that we can quantify the original transcript abundance with high reproducibility and sufficient accuracy using UMIs.

Next, we analyzed the transcriptomes of single cells extracted from the cells of an excised leaf after 0 h and 24 h. We extracted the cell contents from 32 and 34 cells at 0 h and 24 h, respectively, which were used for the preparation of NGS libraries with 1cell-DGE. A total of 2.8 million and 8.5 million reads were obtained, representing mapping rates of 89.9% and 91.5% for the 0 h and 24 h samples, respectively (Additional file 2: Supplementary Figure S5, Additional file 3: Supplementary Table S2). These numbers indicate that our 1cell-DGE method can be used to efficiently construct NGS libraries. Although 98.3% of the mapped reads were removed when the UMI-unifying was performed (Additional file 2: Supplementary Figure S6, Additional file 3: Supplementary Table S2), our simulation of the relationship between read counts and the UMI-unified rate indicated that these read counts are more than sufficient to analyze the transcriptome profiles of the cells used. Our results suggested that a UMI-unified rate of one to two million read counts per sample were sufficient to enable the estimation of the expression profiles to a similar level as that of five million counts per sample, which is similar to estimations reported in previous studies [6] (Additional file 2: Supplementary Figure S7). Our 1cell-DGE approach therefore generated adequate reads for single-cell transcriptome analyses.

### Gene expression profiles of individual leaf cells during reprogramming

We executed the SinQC program [41] to check the library quality of the single-cell samples, as it was not known which mRNAs would be similarly abundant in the individual cells. Based on the distribution of the NGS statistics, all but one of the 66 samples met the quality criteria (Additional file 3: Supplementary Table S2). While SinQC is suitable for the quality control of single-cell transcriptome data without an internal control [41], alternative methods may be more appropriate for the quality control of samples from cell populations including rare cell types or with a smaller number of samples [51].

We identified 2,382 and 4,000 DEGs that were more highly expressed in cells facing the cut edge of a leaf at 0 h and 24 h after excision, respectively (Figures 2, 4). Similar numbers of 0 h-high DEGs were identified in transcriptome profiles generated using whole excised leaves [19, 21], although only 751 genes were found to overlap when using the 5′DGE and 1cell-DGE methods. This may indicate that whole gametophores, comprising a variety of cell types in addition to the leaves, were sampled at 0 h in the study using 5′DGE [19, 21]. On the other hand, a six-fold greater number of DEGs were found to be more highly expressed after 24 h using 1cell-DGE compared to 5′DGE. This result is concordant with the fact that the whole excised leaves at 24 h after excision would have contained more heterogeneous cells, such as reprogramming and non-reprogramming cells, than those at 0 h. The 1cell-DGE approach was highly sensitive to differences in the expression of cell-state specific genes because only the cells facing the cut were analyzed.

The GO term enrichment analysis revealed that the DEGs were enriched in biological process terms related to specific cell states, with photosynthesis genes being more highly expressed at 0 h, while genes involved in the cell cycle, cell differentiation, translation, and DNA metabolic processes were upregulated at 24 h after excision (Figure 3). The expression of *PpCYCD;1,* a partner of *PpCDKA,* which coordinates cell cycle progression and the acquisition of the protonema cell characteristics involved in reprogramming [18], was not detected at 0 h; however, it was detected in many cells at 24 h (Figure 7). Furthermore, *PpCSP1* and *PpCSP2*, which were identified as the common reprogramming factors among plants and animals [20], were more highly expressed at 24 h after the leaf excision than at 0 h (Figure 7). Our results are consistent with previous works related to reprogramming in Physcomitrella, in which low levels of *PpCSP1* promoter activity were detected in the cells of intact leaves, but drastically upregulated in cells facing the cut edge of a leaf [20]. On the other hand, we also detected several cells at 0 h with high levels of *PpCSP1* expression and some at 24 h with low levels of *PpCYCD;1* (Figure 7). These variations most likely reflect the heterogeneity of the cells at the cutting edge, where some cells are reprogrammed into stem cells but others are not [22]. By contrast, the top 10 DEGs detected using 1cell-DGE exhibited no or low levels of expression at 0 h and high levels of expression at 24 h after the leaf excision (Figure 5). These genes may be suitable for use as new cell state markers to discriminate between resting and reprogramming leaf cells in future research.

In addition to these conventional analyses of transcriptomes, pseudotime is an attractive concept for use with scRNA-seq, because the trajectory of the cell states can be predicted even if not all of the various states of the cell profile have been sampled in the analysis [11, 43]. Using only the profiles of individual leaf cells at 0 h and 24 h after the leaf excision, the transcriptome profiles were found to be ordered according to pseudotime (Figure 6b). This suggests that the gene expression profiles at 0 h and 24 h fluctuated and might indicate the pattern of reprogramming in cells facing the cut. We found that pseudotime was correlated to the numbers of detected genes (Additional file 2: Supplementary Figure S9), suggesting that thousands of genes are transiently expressed during reprogramming or that the number of expressed genes increases during the reprogramming of leaf cells into stem cells. Furthermore, at the late phases of pseudotime, the transcriptomes of the cells sampled after 24 h appeared to be separated into two subpopulations with higher and lower numbers of detected genes (Additional file 2: Supplementary Figure S9). This may be the result of the spontaneous arrest or lateral inhibition of the reprogramming of some cells [22]. To clarify this in future research, individual cells separated from other cells [40] and cells facing the cut edge of the leaf should be analyzed at different time points.

## Conclusions

We established 1cell-DGE with microcapillary manipulation as a new scRNA-seq technique, successfully using it to profile the transcriptomes of single cells with high reproducibility and accuracy. Although 1cell-DGE is not a method with as high of a throughput as automated single-cell preparation solutions such as Fluidigm C1 [7] and inDrop [8, 9], it can be used to analyze the contents of single cells from living tissue and organs without the preparation of isolated cells and the associated loss of positional information. This will not only widen the scope of single-cell transcriptome analyses using various types of cells, but also contribute to novel insights into cell–cell interactions in the complicated higher-order structures of multicellular organisms.

## Methods

### Plant materials and growth conditions

The wild-type moss *Physcomitrella patens* Gransden 2004 [38] and the transgenic moss line GX8-NGG [31] were used for the total RNA extractions and the preparation of excised leaves, respectively. To propagate the gametophores, a small portion of GX8-NGG protonema was inoculated on BCDAT agar medium [52] and cultured in a growth chamber (MLR-352H: Panasonic, Tokyo, Japan) under 20–70 μmol/m^2^/s of continuous white light and 55% relative humidity at 23°C.

### Preparation of excised leaves

Gametophores were cultured for 21 days after inoculation on BCDAT medium, after which the distal half of the third leaf was cleanly cut with a razor blade, placed onto the BCDAT medium and covered with cellophane. The majority of the excised leaf, except for the living leaf cells facing the cut edge, was covered with additional layers of cellophane. Dishes containing the excised leaves were sealed with Parafilm and incubated under continuous white light at 23°C until the cell contents were extracted.

### Micromanipulation to extract cell contents

A set of oil hydraulic micromanipulators (MMO-220A and MMO-202ND; Narishige, Tokyo, Japan) and motor-driven manipulators MM-89 (Narishige) were equipped onto an inverted fluorescent microscope IX-70 (Olympus, Tokyo, Japan) with a fluorescence filter unit (U-MWIB3; excitation: 460–495 nm, emission: 510IF; dichromatic mirror: 505 nm; Olympus). To simultaneously observe the tip of the microcapillary and the GFP fluorescence in the nuclei of the leaf cells, the fluorescence microscopy was performed under dim bright-field illumination. The 1.0-mm capillary holder was connected to a microinjector (CellTram vario; Eppendorf, Hamburg, Germany) via a silicone tube filled with mineral oil (M-8410; Sigma-Aldrich, St. Louis, MO, USA), which was in turn attached to the MMO-220A micromanipulator. The parameters of the 1.0-mm glass capillaries were as follows: inner diameter: 20 μm, pipette form: straight, beveled angle of tip: 40°, and pipette length: 55 mm (BioMedical Instruments, Zöllnitz, Germany). The bottom of the glass capillary contained a small amount of cell content extraction mix2 comprising 13% 10× PCR buffer II (Thermo Fisher Scientific, Waltham, MA, USA), 7.8% 25 mM MgCl_2_, 6.5% 0.1 M DTT, 2.6% RNasin Plus RNase inhibitor (Promega, Madison, WI, USA), and 2.6% of a mix containing 2.5 mM of each dNTP (Takara Bio, Kusatsu, Japan). The capillary was attached to the capillary holder so that the beveled tip faced down without any air bubbles.

The attached microcapillary was gently filled with mineral oil under a microscope using the CellTram vario. After adjusting the tip position of the glass capillary to the center of the observation field, a dish containing an excised leaf was set on the microscope and the tip of the capillary was used to extract the nucleus and some surrounding cytoplasm from the target cell. The cell contents were immediately transferred into a 0.2-ml PCR tube containing 1.25 μl RT oligos (0.05 μM) and 2.35 μl cell content extraction mix1, containing 0.45 μl 10× PCR buffer II (Thermo Fisher Scientific), 0.27 μl MgCl_2_ (25 mM), 0.225 μl DTT (0.1 M), 0.09 μl RNasin Plus RNase inhibitor (Promega), 0.09 μl of a mix containing 2.5 mM of each dNTP (Takara Bio), and 0.1 μl of 20,000-fold diluted ERCC RNA spike-in mix (Thermo Fisher Scientific). After a brief centrifugation, the samples were primed with an incubation on a thermal cycler at 70°C for 90 s, 35°C for 15 s, and cooled to 4°C. The tubes were kept on ice before reverse transcription. The RT oligos used in this work are listed in Additional file 3: Supplementary Table S3. Extracted nuclear conditions were categorized into one of six sample quality classes: broken, damaged; broken, average quality; broken, good quality; broken, very good quality; intact, good quality; intact, very good quality.

### Preparation of cDNA libraries for 1cell-DGE

For the reverse transcription, a 0.9 μl RT mix containing 0.33 μl SuperScriptIII reverse transcriptase (Thermo Fisher Scientific), 0.05 μl RNasin plus RNase Inhibitor (Promega), and 0.07 μl T4 gene 32 protein (New England Biolabs, Ipswich, MA, USA) was added to each primed RNA solution. After pipetting gently and centrifuging briefly, the tubes were incubated on a thermal cycler at 50°C for 30 min, 70°C for 10 min, then cooled to 4°C.

To digest the excess RT oligos, the samples were mixed with 0.8 μl nuclease-free water (Qiagen, Hilden, Germany), 0.1 μl 10 × exonuclease I buffer, and 0.1 μl of 20 U/μl exonuclease I (New England Biolabs). After pipetting gently to mix and a brief centrifugation, the tubes were incubated on a thermal cycler using the following conditions: 4°C for 30 s, 37°C for 30 min, 80°C for 20 min with lid heating at 90°C, and cooled to 4°C. The tubes were then transferred onto ice for at least 1 min.

The poly(dA) mix for poly(dA) tailing with RNaseH was as follows: 4.44 μl nuclease-free water (Qiagen), 0.6 μl 10 × PCR buffer II (Thermo Fisher Scientific), 0.36 μl MgCl_2_ (25 mM), 0.18 μl dATP (100 mM) (New England Biolabs), 0.3 μl of 15 U/μl terminal deoxynucleotidyl transferase (Thermo Fisher Scientific), and 0.12 μl of 5 U/μl RNaseH (New England Biolabs). A 6-μl aliquot of this poly(dA) mix was added to each tube after the exonuclease I treatment. After pipetting to mix and a brief centrifugation, the samples were incubated on a thermal cycler using the following conditions: 4°C for 30 s, 37°C for 1.5 min, 70°C for 10 min with lid heating at 80°C, and cooled to 4°C.

For the second-strand synthesis, the following PCR mix1 was prepared: 50.68 μl nuclease-free water (Qiagen), 15.2 μl 5× Q5 reaction buffer with MgCl_2_ (New England Biolabs), 7.6 μl of each dNTP (2.5 mM) (Takara Bio), 0.76 μμl NUP3 primer (100 μM), and 1.76 μl Q5 Hot Start High-Fidelity DNA polymerase (2 U/μl) (New England Biolabs). A 76-μl volume of PCR mix1 was added into each tube after the poly(dA) tailing, pipetted to mix and briefly centrifuged, then the mixtures were divided into 21-μl aliquots which were transferred into four new 0.2-ml PCR tubes. After centrifuging briefly, the tubes were incubated on a thermal cycler in the following conditions: 95°C for 3 min, 98°C for 20 s, 50°C for 2 min, 72°C for 10 min, then cooled to 4°C.

For the cDNA amplification, PCR mix2 was prepared, containing 12.73 μl nuclease-free water (Qiagen), 3.8 μl 5× Q5 reaction buffer with MgCl_2_ (New England Biolabs), 1.9 μl of each dNTP (2.5 mM) (Takara Bio), 0.19 μl BTEP7v2 primer (100 μM), and 0.38 μl Q5 Hot Start High-Fidelity DNA polymerase (2 U/μl) (New England Biolabs). A 19-μl volume of PCR mix2 was added to each tube after the second-strand synthesis, pipetted to mix and briefly centrifuged. The tubes were incubated on a thermal cycler using the following conditions: an initial denaturation at 95°C for 3 min; followed by 22 cycles of 98°C for 10 s, 60°C for 30 s, and 72°C for 6 min, which extended by 6 s at 72°C in each cycle; and stored at 4°C.

After the PCR amplification, the cDNA libraries were purified using a Purelink PCR purification kit with Binding Buffer High-Cutoff (Thermo Fisher Scientific), according to the manufacturer’s instructions. To check availability of each sample, the quantity and quality of the cDNA libraries were measured using a Bioanalyzer 2100 with a High Sensitivity DNA kit (Agilent Technologies, Santa Clara, CA, USA). Each cDNA library solution was placed in a 1.5-ml DNA Lo-bind tube (Eppendorf) and adjusted to a volume of 35 μl with elution buffer (EB) containing 10 mM Tris-HCl (pH 8.0). To remove the byproducts in the cDNA libraries, a 0.55× volume of SPRIselect beads (Beckman Coulter, Brea, CA, USA) were added to each cDNA library solution, which adhered the appropriately sized cDNAs. The tubes were placed on a Magna stand (Nippon Genetics, Tokyo, Japan) for 3 min, and the beads collected at the bottom of the tube. The supernatants were gently removed by aspiration, after which the beads were rinsed twice with 80% ethanol. After air-drying for 10 min, the beads were resuspended in 50 μl EB then left to stand on the Magna stand for 3 min. The resulting supernatants were recovered into new 1.5-ml DNA Lo-bind tubes. Purification with the SPRIselect beads was carried out at least three times. The quantity and quality of the purified cDNA libraries were measured using a Bioanalyzer 2100 with a High Sensitivity DNA kit (Agilent Technologies), and the purified cDNA libraries were stored at –30°C until required. The oligo DNAs used in this work are listed in Additional file 3: Supplementary Table S3.

### Bulk treatment for NGS library construction

In order to fragment the cDNAs to construct the NGS libraries, 2.5 nmol each of four or five purified cDNA libraries were combined, and the volume of the resulting solution was increased to 75 μl with EB. The mixtures were transferred into microTUBE AFA Fiber Pre-Snap-Cap tubes (Covaris, Woburn, MA, USA) and care was taken to prevent any air bubbles. cDNA shearing with a target peak of ~400 bp was carried out using an acoustic solubilizer, Covaris S2 (Covaris), under the following conditions; bath temperature: 4~8 °C, degassing mode: continuous, power mode: frequency sweeping, duty cycle: 10%, intensity: 3, cycles/burst: 200, and time: 90 s. After this treatment, the fragmented cDNAs were transferred to new 1.5-ml DNA Lo-bind tubes (Eppendorf) and purified with a MinElute PCR purification kit (Qiagen). The quality of the fragmented cDNA was measured using a Bioanalyzer 2100 with a High Sensitivity DNA kit (Agilent Technologies).

To recover the fragmented cDNAs tagged with biotin, 20 μl of streptavidin-linked beads, Dynabeads MyOne C1 (Thermo Fisher Scientific), were rinsed twice with 2× BWT buffer containing 10 mM Tris-HCl (pH 7.5), 1 mM EDTA, 2 M NaCl, and 0.02% Tween-20, then suspended in an equal volume of the fragmented cDNA solution. The solutions were left to stand for 10 min to bind the biotinylated cDNA fragments, then placed on a Magna stand for 30 s. The supernatants were discarded and the beads were rinsed three times with 1× BWT buffer and resuspended in 25 μl EBT buffer, which contained 10 mM Tris-HCl (pH 8.5) and 0.02% Tween-20.

For the end repair, a 25-μl mixture containing 5 μl 10× NEBnext End Repair reaction buffer (New England Biolabs) and 2.5 μl of NEBnext End Repair Enzyme Mix (New England Biolabs) was added to the mixture of beads and cDNA fragments. The solution was mixed by gently pipetting and centrifuging briefly, then incubated at room temperature with shaking at 400 rpm for 30 min. The tubes were stood on a Magna stand for 30 s and the supernatants were discarded. The beads were then rinsed twice with EBT buffer while on the Magna stand, after which the stand was removed to enable the beads to be resuspended in 21 μl EBT buffer.

For the dA-tailing, 4 μl of a mixture containing 2.5 μl 10× NEBnext dA-Tailing reaction buffer (New England Biolabs) and 1.5 μl of Klenow Fragment (3′ -> 5′ exo-) (New England Biolabs) was added to the solution of beads with end-repaired cDNA fragments. The mixtures were pipetted gently to mix and briefly centrifuged before being incubated at 37°C with shaking at 400 rpm for 30 min. To remove the reaction mix, the tubes were stood on a Magna stand for 30 s and the supernatants were discarded. The beads were rinsed twice with EBT buffer and resuspended in 25 μl EBT buffer.

To ligate the adapters to the cDNA, a 25-μl mixture containing 5 μl 10× T4 DNA ligase buffer (New England Biolabs), 1.5 μl RP1 adaptor v2 (100 μM), and 5 μl of 400 U/μl T4 DNA ligase (New England Biolabs) was added to the solution containing the beads and dA-tailed cDNA fragments. The solutions were mixed by gently pipetting then centrifuged briefly, after which they were incubated at 20°C with shaking at 400 rpm for 20 min. A 5-μl aliquot of 1 U/μl USER enzyme mix (New England Biolabs) was added to each tube, pipetted gently to mix, then incubated at 37°C with shaking at 400 rpm for 60 min. To remove the reaction mix, the tubes were stood on a Magna stand for 30 s and the supernatants were discarded. The beads were rinsed twice with EBT buffer and resuspended in 25 μl EBT buffer.

To fill the 5′ overhang in the cDNA, a 5-μl mixture containing 3 μl 10× NEB buffer 2 (New England Biolabs), 1 μl of a mix containing 2.5 mM of each dNTP (Takara Bio), and 1 μl of 10 U/μl DNA pol I (New England Biolabs) was added to the solution of adaptor-ligated cDNA fragments and beads. After pipetting gently to mix and centrifuging briefly, the tubes were incubated at 37°C with shaking at 400 rpm for 30 min. To remove the reaction mix, the tubes were stood on a Magna stand for 30 s and the supernatants were discarded. The beads were then rinsed twice with EBT buffer and resuspended in 25 μl EBT buffer.

For the library enrichment, a PCR mix3 was prepared containing 50 μl nuclease-free water (Qiagen), 20 μl 5× KAPAHiFi reaction buffer (KAPA Biosystems, MA, USA), 3 μl a mix containing 10 mM of each dNTP (KAPA Biosystems), 3 μl P5RP1 primer (100 μM), 3 μl EP7v2 primer (100 μM), and 2 μl of 1 U/μl KAPAHiFi Hot Start DNA polymerase (Roche, Basel, Switzerland). The 25-μl mixture of 5′-end-filled cDNAs and the beads was transferred into a new 0.2-ml PCR tube and mixed with 75 μl of PCR mix 3 and centrifuged briefly. The beads were resuspended by pipetting, after which the tubes were immediately set on a thermal cycler and a PCR was performed using the following conditions: an initial denaturation at 95°C for 2 min; followed by 10 cycles of 98°C for 20 s, 63°C for 30 s, and 72°C for 30 s; with a final extension at 72°C for 5 min, after which the samples were stored at 4°C. The enriched libraries were purified with a MinElute PCR purification kit (Qiagen) and eluted with 28 μl EB, according to the manufacturer’s instructions.

The NGS libraries were next subjected to size selection, where fragments measuring 300 bp to 800 bp were extracted from the enriched NGS libraries using BluePippin (Sage Science, Beverly, MA, USA) with a 1.5% dye-free agarose gel cassette and the internal standard R2, according to the manufacturer’s instructions. Size-selected NGS libraries were purified with a MinElute PCR purification kit (Qiagen) and eluted in 28 μl EB, according to the manufacturer’s instruction. The quantity and quality of the NGS libraries were determined using a Bioanalyzer 2100 with a High Sensitivity DNA kit (Agilent Technologies). The oligo DNAs used in this work are listed in Additional file 3: Supplementary Table S3.

### qPCR

Each 20-μl qPCR mixture contained 2 μl of the cDNA templates, LightCycler 480 SYBR Green I Master mix (Roche), and 0.5 μM of each primer, which are listed in Additional file 3: Supplementary Table S3. The qPCRs were performed using a LightCycler 480 (Roche) with the following conditions: 95°C for 8 min; followed by 35 to 50 cycles of 95°C for 10 s, 56°C for 20 s, and 72°C for 15 s. After the amplification cycles, the melting curves were checked to confirm the target validity using the following conditions: 95°C for 10 s, 65°C for 1 min, and heating to 97°C while determining the fluorescence intensity of SYBR Green I five times per 1°C increase. The transcript levels (copy numbers and Cp values) were calculated using standard curves for absolute quantification generated using a dilution series (10, 1, 0.1, 0.01, 0.001, 0.0001, and 0.00001 pg/μl) of the following plasmids: *NGG*: pENTR::*NGG* (5.2 kb), *PpCYCD;1* (AJ428953): pJET::*PpCYCD;1* (4.5 kb), *PpEF1α* (XM_001753007): pJET::*PpEF1α* (4.9 kb), and *PpTUA1* (AB096718): pphb6e07 (4.9 kb). Using the molecular weight of the plasmids, the copy numbers of the transcripts were calculated as follows: weight in Daltons (g/mol) = (bp size of plasmids) (615[Da/bp]). Hence, (g/mol)/Avogadro’s number = g/molecule = copy numbers. This calculation produced copy numbers equivalent to the double-stranded DNA. These experiments were carried out and evaluated using three sets of experimental replicates. Missing values were substituted for a value one-tenth of the minimum value of each transcript level.

### NGS

A 10-μl aliquot of 10 nM sequence libraries consisting of either 32 cells facing the cut edge of the leaf at 0 h after its excision or 34 cells facing the cut at 24 h after leaf excision was denatured and loaded into a lane of the flow cell on a HiSeq1500 sequencer (Illumina, San Diego, CA, USA), following the manufacturer’s instructions. The SBS condition in HighOutput v4 was 126 bp of Read1 and 20 bp of index read. The output data in the BCL files were converted to fastq files of read1 and the index read using bcl2fastq package v1.8.4 (Illumina), and the NGS data were deposited in DDBJ (accessions DRA006455 and DRA006456).

### Demultiplexing and tag counting of 1-cellDGE data

For the sequences with 18 bp of index reads, 8 bp were the multiplex index and 10 bp were the UMI. Therefore, 2 bp of 20 bp of index reads were trimmed from the 3′ end of the reads in the fastq files, using the cutadapt package [53]. To count the numbers of UMIs obtained from the 1cell-DGE data, the fastq files of read1 and the trimmed index read were processed by the package UMI_SC (Nishiyama, 2016: https://github.com/tomoakin/UMI_SC). The 10-bp UMI sequences were inserted into the name of read1, then the sequences were demultiplexed using 6 bp or 8 bp within the trimmed index read for each sample. The read1 sequences were then trimmed and filtered with the trimmomatic package (v0.36) [54] using the options "ILLUMINACLIP:adapters_1cellDGE.fa:2:30:7 TRAILING:20 SLIDINGWINDOW:4:15 MINLEN:30", after which they were mapped onto the reference transcripts using bowtie (v1.1.2). Subsequently, one mapped read was randomly selected from the all mapped reads with the same UMI that are located on the same gene. Finally, the UMI counts for the genes and transcripts (isoforms) were estimated using RSEM (v1.3.0) [55] and exported a table of 1cell-DGE data. The read1 data were mapped on Physcomitrella v3.3 gene models [39] (Phytozome12: https://phytozome.jgi.doe.gov/pz/portal.html).

### Statistical analyses of 1-cell DGE data

To check the quality of 1cell-DGE data, the SinQC program [41] was run using the following settings: MAX FPR: 0.05, TPM Cutoff: 1, Spearman’s test p-value: <0.001, and Pearson’s test p-value: <0.001. The UMI counts were normalized using a scaling normalization method with iDEGES implemented in the TCC package [42], using trimmed mean of M (TMM) values [56] and an exact test of edgeR [57] with the following settings: norm.method = "tmm", test.method = "edger", iteration = 3, FDR = 0.1, and floorPDEG = 0.05.

To detect DEGs, the q.value was set to FDR < 0.01. GO term enrichment analysis was performed using cytoscape v3.4.0 with a BinGO plug-in, and the ontology of biological processes was assessed using GOSlim_plants. The BinGO settings were as follows: statistical test: binomial test, multiple testing correction: Benjamini & Hochberg False Discovery Rate (FDR), and significance level: 0.01. Most of the statistical analyses were performed using R v3.3.3 with Rstudio v0.99.491. The plots were drawn using ggplot2 package v2.2.1. The ICA and pseudotime calculations were carried out using the monocle package v1.4.0 [43]. Hoeffding’s *D* tests of independence were performed using Hmisc package v4.0-3 [44].

## Abbreviations

1cell-DGE: Single-cell digital gene expression
5′DGE: Digital gene expression profiling method using mRNA 5′-end tags
bp: Base pair
cDNA: Complementary DNA
Cp: Closing point
dA: Deoxyadenine
DEG: Differentially expressed gene
dNTP: Deoxyadenosine triphosphate, deoxythimine triphosphate, deoxyguanosine triphosphate, deoxycytidine triphosphate
dT: Deoxythymine
EB: Elution buffer
ERCC: External RNA controls consortium
FDR: False discovery rate
FPR: False positive rate
GFP: Green fluorescent protein
GO: Gene Ontology
GUS: β-glucuronidase
ICA: Independent component analysis
iDEGES: Iterative differentially expressed gene exclusion strategy
NGS: Next generation sequencing
PCR: Polymerase chain reaction
qPCR: Quantitative PCR
RNase: Ribonuclease
RT: Reverse transcription
SBS: Sequencing by synthesis
SE: Single-end
TPM: Transcripts per million
TdT: Terminal deoxynucleotidyl transferase
UMI: Unique molecular identifier

## Declarations

## Acknowledgments

We are grateful to Dr. Olaf Faustmann at Eppendorf for help with the micromanipulation. We also thank Drs. Shuji Shigenobu and Katsushi Yamaguchi at the National Institute for Basic Biology (NIBB) and Drs. Asao Fujiyama and Atsushi Toyoda at the National Institute of Genetics for their help with the next-generation sequencing. We are grateful to Dr. Yukiko Kabeya at NIBB and Ms. Ritsuko Okamoto at the Nara Institute of Science and Technology for their technical assistance. The computational analyses were partially performed on the NIG supercomputer at the ROIS National Institute of Genetics.

## Authors’ contributions

MK, RR, and MH conceived and designed the experiments. MK, TN, YT, MI, TM, and AI established the experimental conditions. MK prepared the plant materials, performed the micromanipulation, constructed the cDNA and NGS libraries, and performed the qPCR. MK, TN, RS, YT, and DL carried out the computational analyses of the NGS data. TN developed the UMI_SC package. MK, RR, TN, and MH wrote the paper. YT, MI, TM, AI, and TD revised the manuscript. All authors read and approved the final manuscript.

## Ethics approval and consent to participate

Not applicable.

## Consent for publication

Not applicable.

## Availability of data and material

The wild-type moss *Physcomitrella patens* Gransden 2004 [38] and the transgenic moss line GX8-NGG [31], as well as the plasmids pENTR::*NGG*, *PpCYCD;1* (AJ428953), *PpEF1α* (XM_001753007), and *PpTUA1* (AB096718); pphb6e07 are available from the corresponding author Mitsuyasu Hasebe at the National Institute for Basic Biology (NIBB), Japan. The package UMI_SC developed by TN is available on GitHub (https://github.com/tomoakin/UMI_SC). The datasets from the next-generation sequencing performed in this study are available in DDBJ (accession DRA006455, DRA006456).

## Publisher’s Note

Springer Nature remains neutral with regard to the jurisdictional claims in published maps and institutional affiliations.

## Additional files

Additional file 1: ***Supplementary Movie S1.*** Extraction of the contents of a single cell in an excised leaf of Physcomitrella using a microcapillary. GFP fluorescence was observed using the fluorescence filter unit (see Methods) under a mercury arc lamp and with dim bright-field illumination.

Additional file 2: Supplementary Figures S1 to S9.

***Supplementary Figure S1.*** Transcript levels of *NGG*, *PpEF1α*, *PpCYCD;1*, and *PpTUA1* in the contents of single cells.

***Supplementary Figure S2.*** Abundance of *PpTUA1* cDNA in the samples reverse-transcribed from different concentrations of total RNA.

***Supplementary Figure S3.*** Validation of 1cell-DGE by pilot sequencing.

***Supplementary Figure S4.*** Schematic representation of the 1cell-DGE workflow

***Supplementary Figure S5.*** Statistics of 1cell-DGE performed on single leaf cells from Physcomitrella.

***Supplementary Figure S6.*** Quality check of 1cell-DGE data.

***Supplementary Figure S7.*** Estimation of sequencing coverage using 1cell-DGE.

***Supplementary Figure S8.*** Independent component analyses of 1cell-DGE data.

***Supplementary Figure S9.*** Correlation between pseudotime and NGS statistics.

Additional file 3: Supplementary Tables S1 to S3.

***Supplementary Table S1.*** Statistics of pilot sequencing for 1cell-DGE.

***Supplementary Table S2.*** NGS statistics of 1cell-DGE data derived from individual leaf cells at 0 h and 24 h after leaf excision.

***Supplementary Table S3.*** Oligo DNAs used in this work.

**Supplementary figure legends**

***Supplementary Figure S1. Transcript levels of* NGG, PpEF1α, PpCYCD;1*, and* PpTUA1 *in the contents of single cells.*** (a) Dot plots show the mean number of transcripts converted to cDNA using reverse transcription, derived from technical triplicates measured using qPCR. The numbers along the x axis represent 10 independent samples derived from the contents of single cells. Error bars indicate the standard deviation derived from technical triplicates. (b) Jitter plots show the mean Cp values of the transcripts. Each dot represents an independent single cell sample.

***Supplementary Figure S2. Abundance of* PpTUA1 *cDNA in the samples reverse-transcribed from different concentrations of total RNA.*** The jitter plot shows the Cp values as the abundance of *PpTUA1* cDNA derived from different concentrations of total RNA. The cDNA was generated with either poly(dA) tailing or template switching.

***Supplementary Figure S3. Validation of 1cell-DGE by pilot sequencing.*** Scatter plots of the expression profiles of Physcomitrella genes (a, b) or the ERCC RNA spike-in mix (c–h) derived from 5 μg and 20 pg of total RNA. The 20 pg total RNA was diluted from 5 μg of total RNA including a 0.1-fold ERCC RNA spike-in mix with nuclease-free water. Illumina sequencing libraries of replicate 1 (Rep1) and replicate 2 (Rep2) were prepared from the same RNA stock solution. For the 5 μg of samples, the cDNA amplification step was skipped when generating the 1-cell cDNA libraries (see Supplementary Figure S4, Methods) was skipped. (a, b, c, d) Rep1 and Rep2 of total read (grey) and UMI counts (black) in which the unified duplicated reads with the same UMI mapped on the same gene locus were plotted. (e, f, g, h) UMI counts of the ERCC RNA spike-in mix in each replicate were plotted against the concentration of the ERCC RNA spike-in mix. R^2^ indicates the determination coefficients.

***Supplementary Figure S4. Schematic representation of the 1cell-DGE workflow.*** There are four major steps to 1cell-DGE: 1. extraction of the contents of a single cell using microcapillary manipulation; 2. preparation of 1-cell cDNA libraries using reverse transcription (RT) with RIU (RP2 sequence for index read priming, index sequence, and unique molecular identifier (UMI)) RT oligos; 3. bulk treatment of Illumina next-generation sequencing (NGS) libraries; and 4. NGS is performed. NGS was performed on Illumina sequencers using the following conditions: single-end read (SE): 50 or 126 bp, index read: 18 bp, including 8 bp of multiplex index and 8 or 10 bp of random oligonucleotide sequence for the UMI. B indicates a biotin modification at the 5′ end of the DNA oligonucleotides for cDNA amplification, which enables the capture of the cDNA fragments using avidin-conjugated magnetic beads.

***Supplementary Figure S5. Statistics of 1cell-DGE performed on single leaf cells from Physcomitrella.*** Box and jitter plots indicate the statistics of the 1cell-DGE data generated for 32 and 34 individual cells sampled at 0 h and 24 h after leaf excision. These read counts include the reads mapped onto the Physcomitrella v.3.3 gene models [39] and the ERCC RNA spike-in mix before their unification using the UMIs. (a) Total read counts. (b) Mapped read counts. (c) Mapping rate (Additional file 3: Supplementary Table S2).

***Supplementary Figure S6. Quality check of 1cell-DGE data.*** Box and jitter plots show (a) the total number of UMI counts, (b) the UMI-unified rate, (c) the number of detected genes, and (d) the read complexity of the 32 and 34 individual cells sampled at 0 h and 24 h after leaf excision, respectively (Additional file 3: Supplementary Table S2). These statistics were calculated using the SinQC package [41] and the following settings: Max FPR: 0.05, TPM Cutoff: 1, Spearman’s test p-value: <0.001, and Pearson’s test p-value: <0.001.

***Supplementary Figure S7. Estimation of sequencing coverage using 1cell-DGE.*** The relationships between the sequencing depth and (a, b) the number of detected genes and (c, d) the UMI-unified rate are presented. 0.1, 0.2, 0.5, 1.0, 2.0, and 5.0 million reads were randomly sampled from 1cell-DGE data derived from the individual leaf cells Pp0h07, Pp0h08, Pp0h42 at 0 h after excision (a, c) and Pp24h07, Pp24h08, Pp24h42 at 24 h after excision (b, d). These sample data were processed using the UMI_SC package (Nishiyama, 2016: https://github.com/tomoakin/UMI_SC) and were counted using the SinQC program [41]. Triangles represent Pp0h07 and Pp24h07 values with the index sequence ATACGTGC. Squares represent Pp0h08 and Pp24h08 values with the index sequence CGTGCATA. Circles represent Pp0h42 and Pp24h42 values with the index sequence ACTTCGGT.

***Supplementary Figure S8. Independent component analyses of 1cell-DGE data.*** 1cell-DGE data were calculated to reduce the dimensions of the expression profiles using an independent component analysis and plotted using the monocle v1.4 package [43]. Each dot indicates independent cell samples which were colored by (a) their extracted nuclear condition, (b) their leaf excision date, (c) the byproduct contamination, and (d) the cDNA amounts. Circles represent cells sampled 0 h after leaf excision, triangles represent cells sampled 24 h after leaf excision.

***Supplementary Figure S9. Correlation between pseudotime and NGS statistics.*** The pseudotimes of the 1cell-DGE data were plotted against (a) the mapped read counts and (b) the numbers of detected genes. Each dot indicates an independent cell categorized by the time it was sampled after the leaf excision. *D* indicates the statistical values calculated using Hoeffding’s *D* test of independence in the Hmisc package [44].

